# Drosophila larval brain neoplasms present tumour-type dependent genome instability

**DOI:** 10.1101/192492

**Authors:** Fabrizio Rossi, Camille Stephan-Otto Attolini, Jose Luis Mosquera, Cayetano Gonzalez

**Affiliations:** Institute for Research in Biomedicine (IRB Barcelona), The Barcelona Institute of Science and Technology, Baldiri Reixac, 10, 08028 Barcelona, Spain.; Catalan Institution for Research and Advanced Studies (ICREA), Barcelona, Spain.

**Keywords:** drosophila cancer model, copy-number variation, single nucleotide polymorphism, genome instability, somatic mutation

## Abstract

Single nucleotide polymorphisms (SNPs) and copy number variants (CNVs) are found at different rates in human cancer. To determine if these genetic lesions appear in Drosophila tumours we have sequenced the genomes of 17 malignant neoplasms caused by mutations *in l(3)mbt, brat, aurA*, or *lgl*. We have found CNVs and SNPs in all the tumours. Tumour-linked CNVs range between 11 and 80 per sample, affecting between 92 and 1546 coding sequences. CNVs are in average less frequent in *l(3)mbt* than in *brat* lines. Nearly half of the CNVs fall within the 10 to 100Kb range, all tumour samples contain CNVs larger that 100 Kb and some have CNVs larger than 1Mb. The rates of tumour-linked SNPs change more than 20-fold depending on the tumour type: late stage *brat*, *l(3)mbt*, and *aurA* and lgl lines present median values of SNPs/Mb of exome of 0.16, 0.48, and 3.6, respectively. Higher SNP rates are mostly accounted for by C>A transversions, which likely reflect enhanced oxidative stress conditions in the affected tumours. Both CNVs and SNPs turn over rapidly. We found no evidence for selection of a gene signature affected by CNVs or SNPs in the cohort. Altogether, our results show that the rates of CNVs and SNPs, as well as the distribution of CNV sizes in this cohort of Drosophila tumours are well within the range of those reported for human cancer. Genome instability is therefore inherent to Drosophila malignant neoplastic growth at a variable extent that is tumour type dependent.

**AUTHOR SUMMARY:** Drosophila models of malignant growth can help to understand the molecular mechanisms of malignancy. These models are known to exhibit some of the hallmarks of cancer like sustained growth, immortality, metabolic reprogramming, and others. However, it is currently unclear if these fly models are affected by genome instability, which is another hallmark of many human malignant tumours. To address this issue we have sequenced and analysed the genomes of a cohort of seventeen fly tumour samples. We have found that genome instability is a common trait of Drosophila malignant tumours, which occurs at an extent that is tumour-type dependent, at rates that are similar to those of different human cancers.

## INTRODUCTION

A wide range of tumour types can be experimentally induced in different organs in Drosophila melanogaster [1-6]. Many of these tumours are hyperplasias that present during larval development and eventually differentiate, but others behave as frankly malignant neoplasms that are refractory to differentiation signals, lethal to the host and immortal. The latter can be maintained through successive rounds of allograft in adult flies [7].

In humans, the study of mutational landscapes in thousands of tumours has generated a large catalogue of genomic lesions that appear during tumour development and are a driving force for malignant growth in different cancer types [8-13]. In Drosophila, the sequencing of a single tumour caused by the loss of Polyhomeotic (Ph) revealed that neither single nucleotide polymorphisms (SNPs) nor copy number variations (CNVs) were significantly increased in comparison with non-tumoural control tissue, suggesting that genome instability (GI) may not be a pre-requisite for neoplastic epithelial growth in this model system. [14]. The question remains, however, as to the extent of GI in other samples of Ph tumours and, indeed, in different types of Drosophila malignant neoplasms.

To address this question we have investigated the mutational landscape of a cohort of tumours caused by mutations in l(3)malignant brain tumour (*l(3)mbt*), *brain tumour* (*brat*), *aurora-A* (*aurA*), and *l(2)giant larvae* (*lgl*), which are some of the most aggressive and best characterised larval brain tumours that can be induced in Drosophila [15-20]. Although similar in appearance under the dissection microscope, these tumours develop through different oncogenic pathways and originate from different cell types. Mutants in *brat*, *aurA*, and *lgl* disrupt different aspects of the mechanisms of neuroblasts asymmetric division. The cell-of-origin of tumours caused by mutation in *brat* tumours is only the type II neuroblast, which resides in the dorsal side of the central brain [17], while *aurA* and *lgl* tumours originate from type I and II neuroblasts [16, 18-20]. Neoplastic growth in *l(3)mbt* tumours originate in the neuroepithelial regions of the larval brain lobes [19, 21] and is tightly linked to the ectopic expression in the soma of germline genes [22].

Altogether, we sequenced a total of 17 genomes corresponding to a combination of tumour types, lines of the same tumour type, lines from the same individual, and time points. Our results show that CNVs and SNPs appear in Drosophila malignant neoplasms at a rate that is tumour-type dependent and within the range reported for human cancer.

## RESULTS/DISCUSSION

To determine the extent of genome instability (GI) in Drosophila malignant neoplasms we generated a cohort from six different larval brain tumours including two *l(3)mbt* (mbtL1 and mbtL2), two *brat* (bra*t*L1 and bratL2), one *aurA*, and one *lgl* (Fig. 1). Following allografts into adult hosts [7], gDNA samples were taken at T0 (first round of allograft), T5, and in some cases T10. One of the *l(3)mbt* tumour lines was split at T9 into two sub-lines that were cultured separately up to T10.

**Fig. 1.**
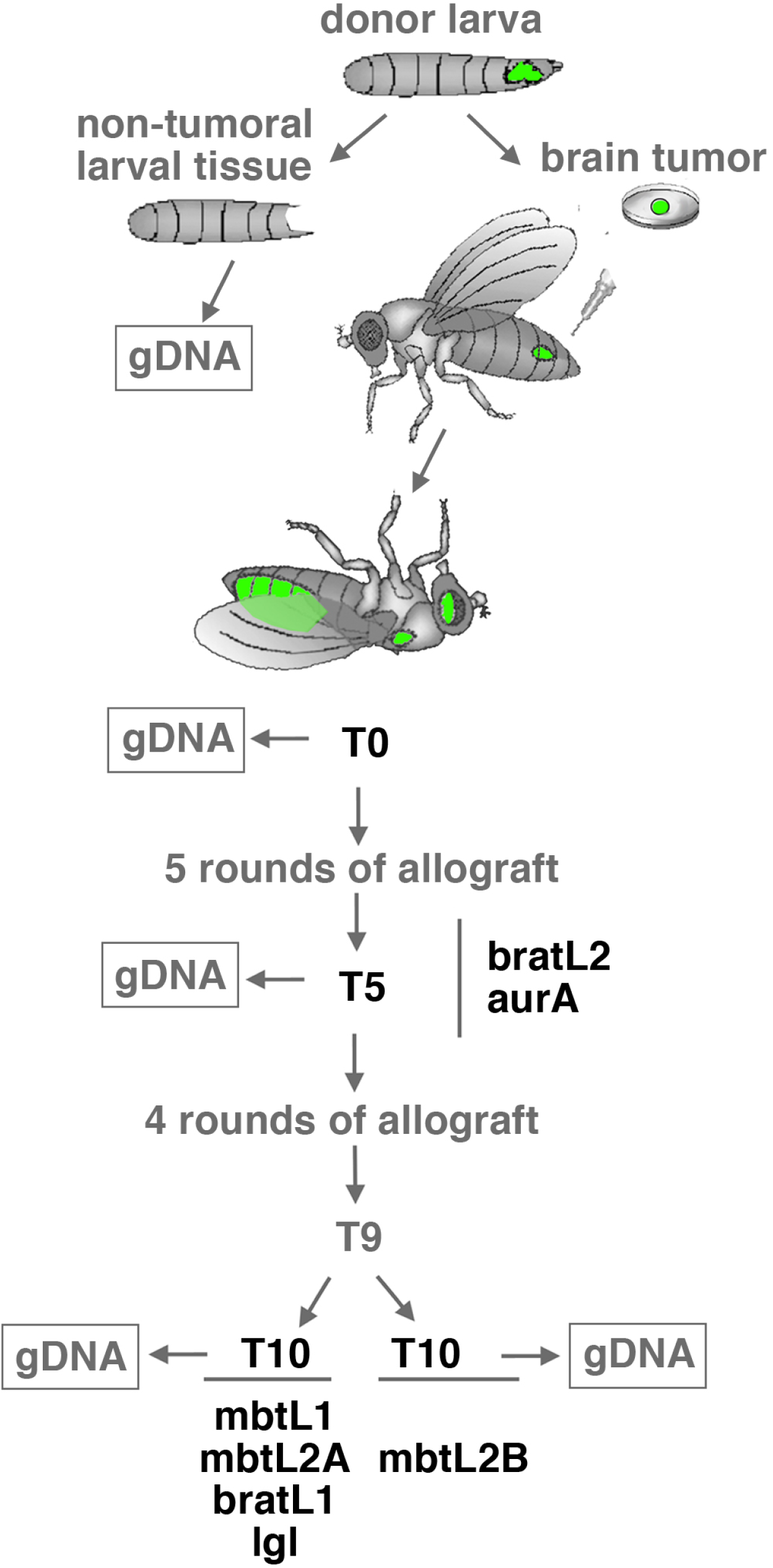
The Drosophila larval brain tumour cohort. Larval brains tumours derived from two *l(3)mbt* (mbtL1 and mbtL2), two *brat* (bratL1 and bratL2) one *aurA*, and one *lgl* individuals were dissected out from the donor larvae and allografted repeatedly, up to T5 for bratL2 and *aurA*, and up to T10 for mbtL1, mbtL2, bratL1, and *lgl*. Line mbtL2 was split at T9 to generate sublines mbtL2A and mbtL2B. Genomic DNA was obtained from all tumour lines at T0, T5, and T10 if available, as well as from the non-tumoural tissues of the corresponding donor larvae.

### CNVs are frequent in the Drosophila brain tumour cohort

To identify copy number variants (CNVs) that appear during tumour growth we compared the gDNA coverage from each tumour sample of the cohort to that of the larva in which each of these tumours originated. Based on the detection of Y chromosome-specific sequences and the ratio of X chromosome / autosomes coverage we concluded that that mbtL1, mbtL2, and *lgl* tumour lines originated from male larvae while aur, bratL1, and bratL2 originated from females (Fig. S1A). We could not sex the tumours before allografting because testis do not develop in some of these mutant larvae. Most (88%) of the identified CNVs correspond to gains clustered on heterochromatic and under-replicated euchromatic regions (URs), which are present in all lines from T0. These regions do not endoreplicate to the full extent that most of the genomic DNA does in polytene larval tissues [23-26] and therefore appear as copy number gains when the non-polytene tumour samples are compared to larval gDNA (Fig. S1B). Their detection provides a valuable internal control for our CNV calling method. Running the algorithm after filtering these regions out with a repeat mask generates the map revealing the actual extent of CNVs that arise during tumour development in our cohort (Table S1). A graphic summary of the map of gains (≥ +2, blue; +1,green) and losses (-1,red; -2, magenta) on each chromosome arm is shown in Fig. 2A. This final filtered map is not only a clean version of the unfiltered; it also includes new CNVs that can only be identified thanks to the finer calibration of diploidy achieved by the algorithm following the removal of URs and heterochromatin.

**Fig. 2.**
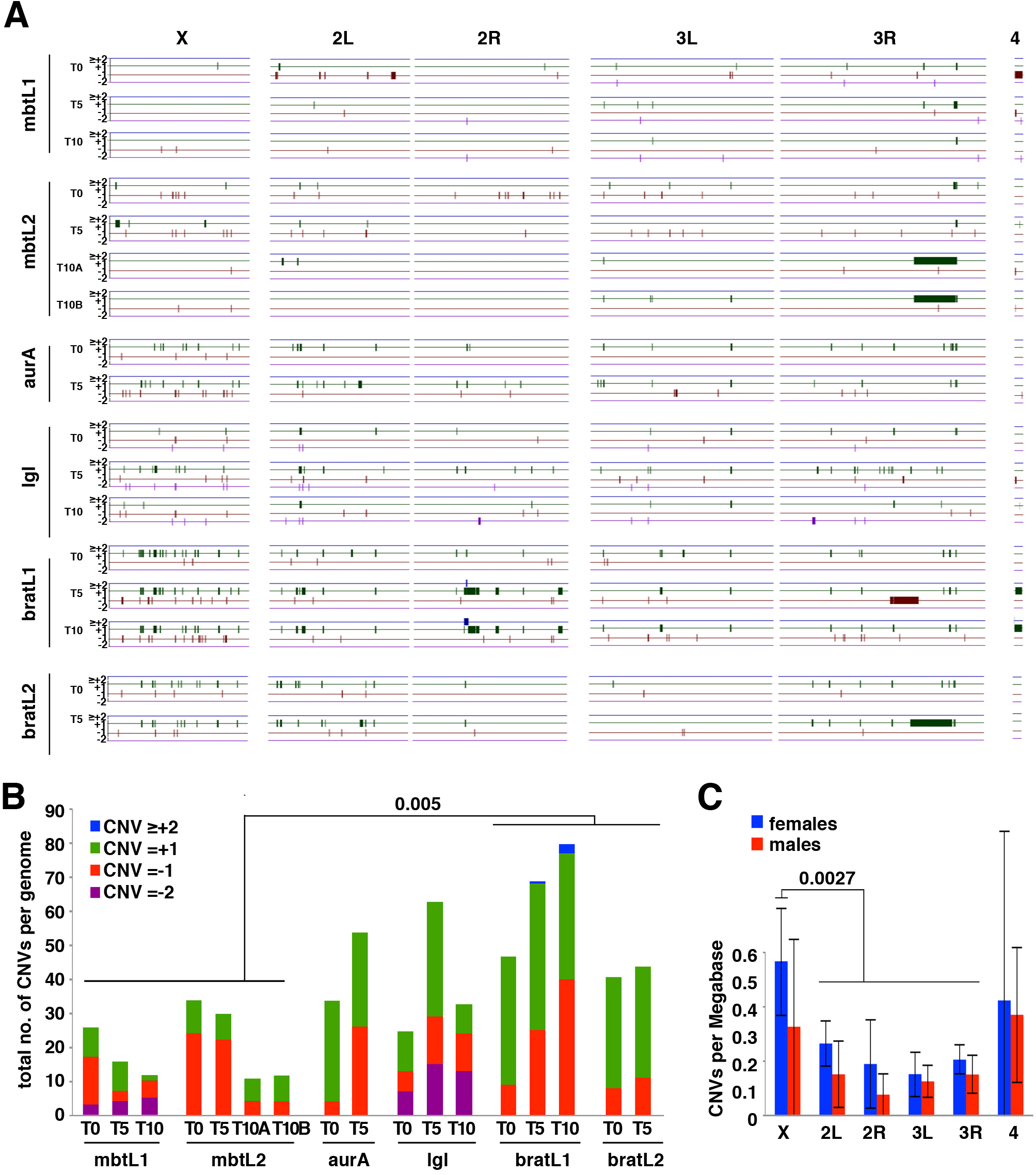
Map and frequency of CNVs. A) Map of the CNVs identified in different lines at different time points after filtering out under-replicated regions. Gains (≥+2, blue; +1, green) and losses (-1, red; and ≤-2, purple) are mapped along chromosome arms X, 2L, 2R, 3L, 3R, and 4th. The heterochromatic Y chromosome is omitted.B) Barplot showing the total number of CNVs per genome per tumour sample and the relative contribution of each of the four CNV classes. C) Distribution of CNVs per Mb on each chromosome arm in female (blue) and male (red) samples. Error bars represent standard deviation.

We detected CNVs in all tumour samples at rates that range between 11 in mbtL2 T10A to 80 in bratL1 T10 (Fig. 2A, B) with an average of 37±20.5. Differences among tumour types are not major, but CNVs per genome are in average significantly fewer in *l(3)mbt* (20.1±9.6) than in *brat* (56.2±17.3; p=0.005) lines. The average number of each CNVs class (-2, -1, +1, and ≥+2) per genome in the entire cohort is 2.8±4.8, 13.5±10.5, 20.6±14, and 0.2±0.7, respectively. The only cases of ≥+2 were observed in the bratL1 line. Gains and losses of a single copy (Fig. 2B, classes +1 and -1; green and red, respectively) account for 92% of the found CNVs, with class +1 being more frequent in 65% or the samples. Amplifications are 1.3 times more abundant than deletions (354 and 277, respectively).

The four largest CNVs found in the cohort, much larger than all the rest, are one deletion and three duplications that, remarkably, fall in the same subdistal region in 3R and overlap extensively. The largest duplication was found in bratL2 T5 and spans 6.9 Mb on chromosome 3 (chr3R:20994001-27965000). This region (Fig. 2A, longest thick green segment) overlaps extensively with two adjacent duplications of 4.0Mb (chr3R:21317001-2539800) and 2.5 Mb (chr3R:25402001-27960000) that are found in both mbtL2 T10A and mbtL2 T10B. Owed to the low resolution of Fig. 2A, the two adjacent duplications appear as a single thick green segment in each tumour line. The large duplication in bratL2 T5 referred to above also overlaps over 1.1 Mb with the 4.1 Mb deletion (chr3R:17979001-22092000) observed in bratL1 T5, the largest deletion found in the cohort.

The rate of CNVs/Mb is slightly smaller in all chromosomes in male (range=0.08-0.37 CNVs/Mb) than in female (range= 0.15-0.57) tumour samples, but differences are poorly significant (Fig. 2C; p=0.055).There are no significant differences in the rate of CNVs per Mb of euchromatin among chromosomes, except for the X chromosome in female samples (0.57±0.2 CNVs/Mb) which is significantly higher than in the autosomes (p=0.0027).

All tumour samples in the cohort present a nearly diploid balance of chromosome stoichiometry (i.e 1X, 1Y, 2A in males; 2X, 2A in females). Most of the Y chromosome cannot be quantified due to the abundance of low complexity sequences and transposable elements (TEs). However, in all tumour samples derived from male larvae the coverage of the repeat-free kl-2 gene region is very close to half of the mean coverage of the major autosomes, regardless of the stage of tumour growth. This result strongly suggests that unlike male cell lines, which often loose the entire Y [27], this chromosome is efficiently maintained in Drosophila tumours. The Y chromosome encodes only a handful of genes, all of them male fertility factors with no known function in the soma and, indeed, X/0 males are viable. However, the Y chromosome heterochromatin has a major impact on epigenetic variation and in modulating the expression of biologically relevant phenotypic variation [28]. Similarly, unlike Drosophila cell lines where widespread loss or gain of the entire chromosome 4 has been reported [27], we have only observed three cases of large segmental aneuplodies for this chromosome in our entire cohort: a deletion (-1) uncovering 93% of the euchromatin and two duplications (+1) covering 79% and 91% of the euchromatin of chromosome 4, respectively. As in many types of human cancer, karyotype changes have been observed in allografts from various larval brain tumours [29]. In flies, these changes do not appear to be sufficient to drive tumourigenesis [30, 31], but it is not known if they are involved in tumour progression. Our results suggest that specific aneuploid combinations are not selected during tumour progression.

### CNVs in tumour samples are larger than those found in Drosophila cell lines, and turn over rapidly

CNV size distribution is highly skewed and notably different between duplications and deletions (Fig. 3A). Nearly half of the CNVs (49% of duplications and 47% of deficiencies) fall within the 10 to 100Kb range, but for those <10Kb, deletions and duplications account for 47% and 12% of the total, while in the >100Kb range the corresponding figures are 6% and 39% respectively. Indeed, most (85%, n=20) of the largest CNVs (≥500Kb) are amplifications that appeared at or after T5 (Table S1).

**Fig. 3.**
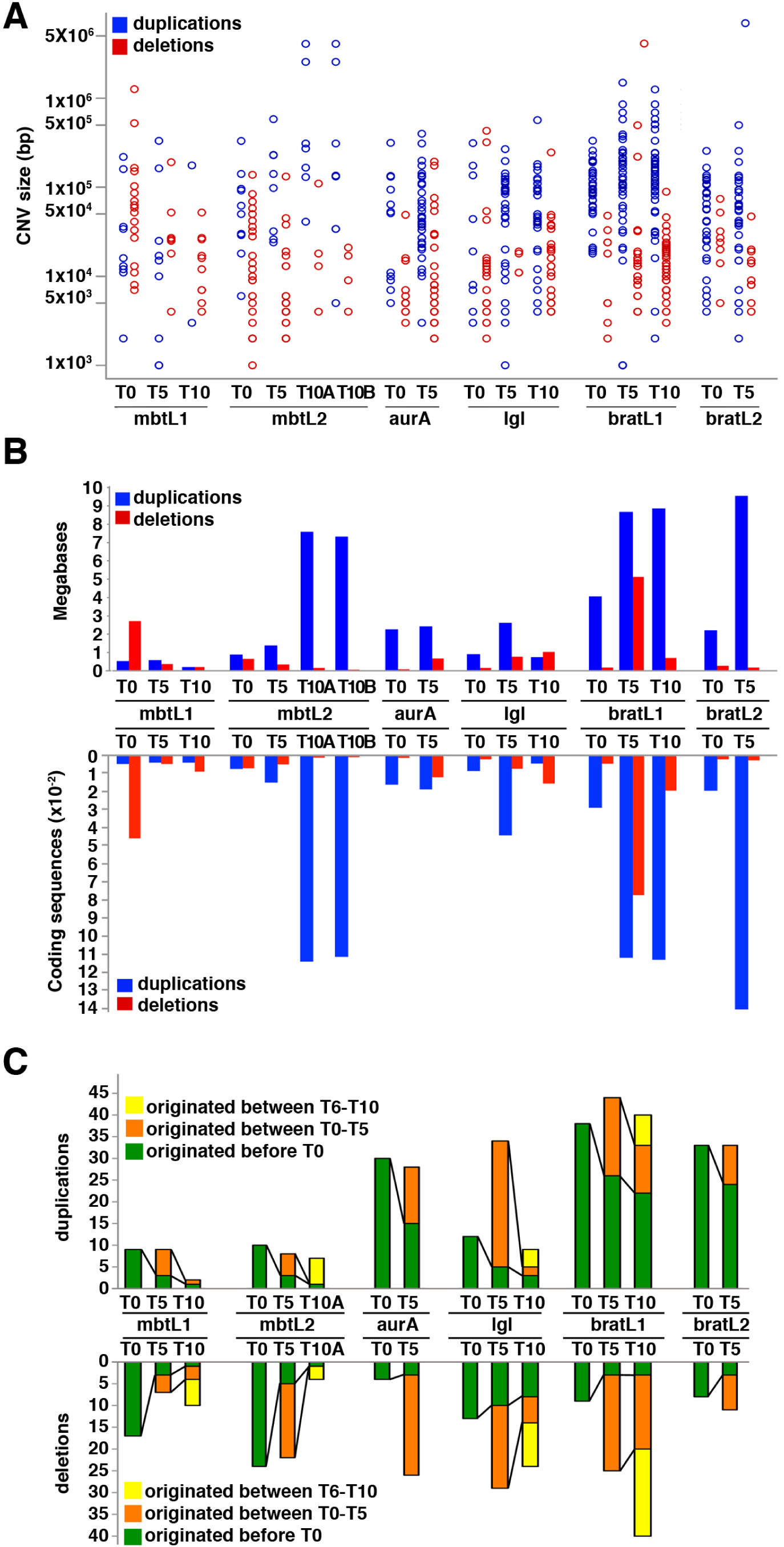
Size distribution and turnover rate of CNVs. A, B) Distribution of CNVs sizes among the samples of the cohort. Duplications and deletions are shown in blue and red, respectively. A shows a scattered plot of CNV sizes in base-pairs, in logarithmic scale. B shows the total number of Mb (upper side of the graph) and total coding sequences (lower side of the graph) affected by duplications and deletions. C) Plot of number of duplications (upper side of the graph) and deletions (lower side of the graph) that are passed on through successive rounds of allograft.

The total length of genomic sequences affected by gains in each tumour sample is quite significant, ranging between 180 Kb and 9.5 Mb. All but one of the 17 samples are affected by duplications covering more than 0.5 Mb. Deletions cover smaller, but still significant regions ranging from 60 Kb to 5.1 Mb. 15 out of 17 samples present deletions covering more than 100 Kb (Fig. 3B). Genomic sequence length correlates tightly with the number of coding sequences affected by copy number variation (Fig. 3B). In the entire cohort the number of genes affected by duplications or deletions range from 40 to 1404 and 9 to 773, respectively. In 11 out of the total 17 samples, duplications affect more than 100 genes and deletions affect more than 30 (Fig. 3B).

Enrichment analysis of the genes duplicated in at least one sample and not deleted in any, shows only proteinaceous extracellular matrix (GO:0005578) as significantly overrepresented, and no GO term was found to be under-represented (Table S2). Proteinaceous extracellular matrix is part of the GO term extracellular region (GO:0005576) that was found to be overrepresented in wild type strains [32]. Enrichment analysis of the genes deleted in at least one sample and not duplicated in any shows that the terms nucleosome assembly (GO:0006334), nuclear nucleosome (GO:0000788), and DNA-templated transcription initiation (GO:0006352), are significantly overrepresented, and no GO term was found to be under-represented (Table S2). However, "nuclear function”, which includes nuclear nucleosome and nucleosome assembly was found to be under-represented in duplicated fragments in wild type strains [32].

The range of CNV sizes found in the tumour cohort is similar to those reported in Drosophila cell lines, and much larger than those found in wild type natural population and laboratory-adapted strains where 95% of the variants are shorter than 5 Kb and the largest duplicated and deleted regions are only 12 kb and 33 kb long, respectively [32-35]. Moreover, unlike Drosophila strains where CNVs affect more frequently regions that do not contain coding sequences [32] [33], 97% of the CNVs found in our tumour cohort affect coding sequences. The range of CNVs length in our tumour cohort is also much larger than those found in a Drosophila epithelial tumour caused by the loss of polyhomeotic (ph) [14] and similar to the 0.5 kb – 85 Mb range found in human cancer [36].

To get an estimate of the rate of turnover of CNVs, we plotted those that appear at any given T together with those that overlap in at least 1 Kb with CNVs found at the previous time point (Fig. 3C). New variants, both amplifications and deletions, appear at each time point, but are diluted at a greater or lesser extent at later stages of tumour growth: the fraction of duplication and deletions passed on from T0 to T10 is within the 5 to 70% range, with no major differences between deficiencies and duplications. More than a third of the total number of CNVs found at any given T were not present at earlier time points. An interesting case reflecting the rate of CNV turnover is that of the pair mbtL2 T10A and T10B. These two samples, which were originated by splitting the mbtL2 line at T9, contained 11 and 12 CNVs respectively of which 8 were common to both lines, thus illustrating a case in which CNVs arise in a single round of transplantation. In total, deficiencies and duplications inherited from T0 account for 7 and 14% of those present at the last round of allograft, respectively.

Three main conclusions can be derived from our results. Firstly, compared to those reported in Drosophila wild type strains, CNVs in our tumour cohort are much more abundant and larger and appear much faster, over a period of weeks rather than years. Such a high rate of interstitial aneuploidy strongly suggests that one or more of the pathways that prevent the formation of interstitial aneuploidies are significantly compromised in these tumours, more in *brat* than in *l(3)mbt*. Secondly, neither number nor size distribution appear to correlate with the stage of tumour growth. This observation strongly argues that the cause of the GI that originates CNVs is concomitant with the onset of neoplastic malignant growth. Finally, their rather random distribution among tumour types and rounds of allografting, rapid turn over, and absence of hotspots shared among different lines suggest that CNVs behave like passengers rather than drivers in these tumours.

### SNPs rates are tumour-type and tumour-age dependent

We used MuTect to call somatic nucleotide polymorphisms (SNPs) between each tumour sample and the non-tumoural tissues of the corresponding larvae (Fig. 4A; Table S3). SNPs in TEs or low complexity sequences were not taken into consideration for further analysis. We found SNPs in all tumour samples, at rates that are tumour type and tumour age-dependent. Total SNP numbers at T0 range between 27 and 76 among all tumour lines and remain unchanged at later time points in the two *brat* lines (range=26-57). However, SNP burden increases to a range between 95 and 218 in the *l(3)mbt* lines and even more, up to 8-fold compared to T0, in the *aurA* and *lgl* lines (range=385-476)(Fig. 4B). A previous report carried out by comparing tumour and control tissue to the Drosophila reference genome found no evidence of tumour-linked SNPs in one sample of allografted Ph tumour at T4 [14]. Using our own SNP calling strategy to directly compare the published tumour and control gDNA sequence we identify 20 tumour-linked SNPs, which is similar to the rate that we have found in the *brat* lines, the ones with the smallest number of SNPs within our cohort.

**Fig. 4.**
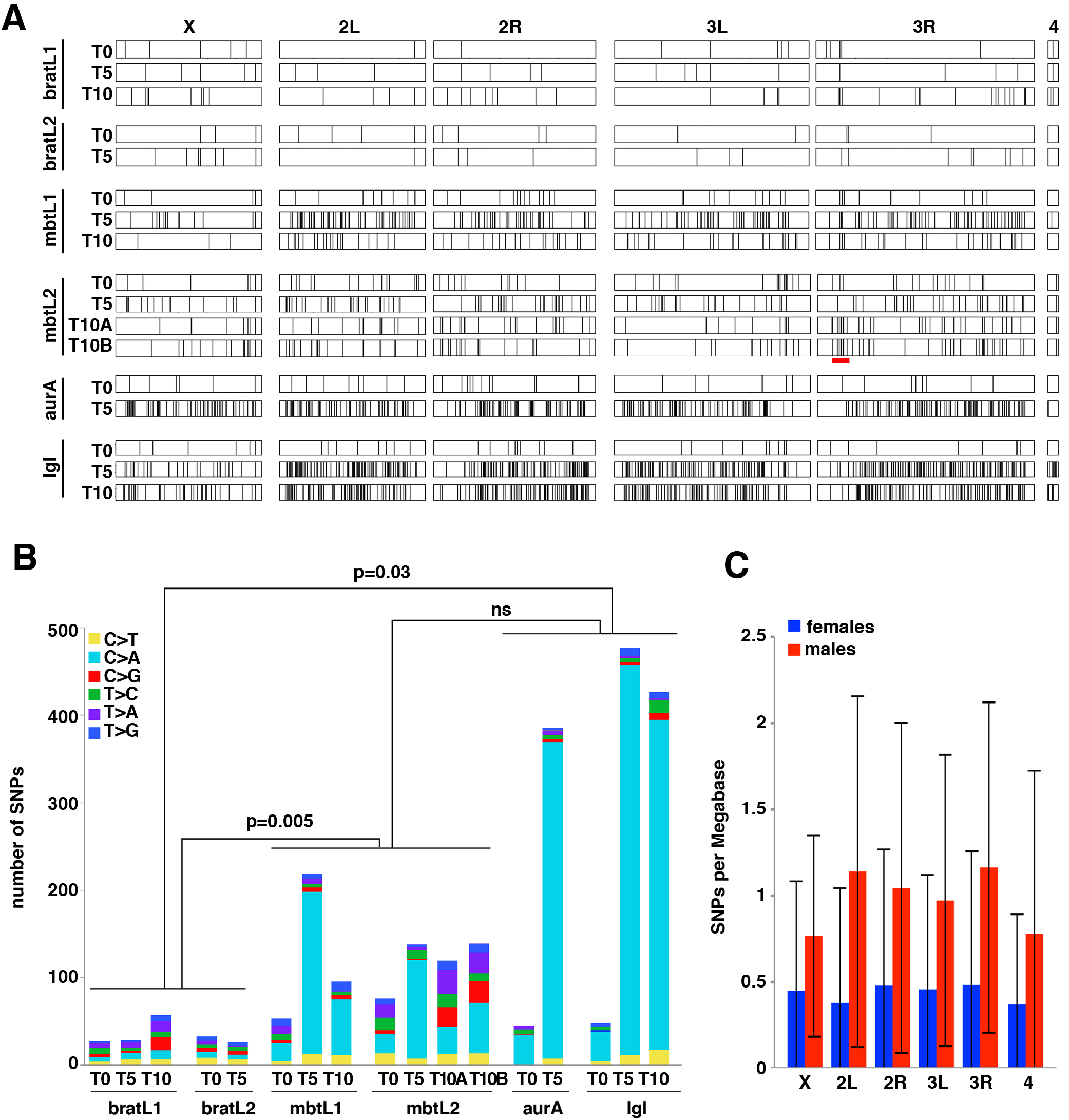
Map and frequency of SNPs. A) The SNPs identified in different lines at different time points are mapped along chromosome arms X, 2L, 2R, 3L, 3R, and 4th. The heterochromatic Y chromosome is omitted.B) Barplot showing the total number of SNPs per genome per tumour sample and the relative contribution of each of the six possible base-pair substitutions. C) Distribution of SNPs per Mb on each chromosome arm in female (blue) and male (red) samples. Error bars represent standard deviation.

Most of the differences in the total number of SNPs among the tumour samples of our cohort are accounted for by C>A (G>T) transversions (Fig. 4B, pale blue) to the extent that such differences among tumour lines at late time points become not significant if these two types of SNPs are removed. Indeed, the increase of C>A transversions becomes particularly notorious at later time points in *aurA* and *lgl* tumour lines where they account for more than 88% of all SNPs (Table S3). Importantly, applying the method described by Costello et al. [37], we were able to discard the possible artifactual origin (i.e. DNA oxidation during the processing of the DNA samples) of the C>A mutations that we have observed. C>A (G>T) transversions are commonly produced by the formation of apurinic (abasic) sites or 8-hydroxy-2′-deoxyguanosine (8-oxo-dG) that result from superoxide anions reacting with deoxyguanosine [38] [39]. Because our sequencing data shows no evidence of mutants in genes involved in the removal of superoxide anions or 8-oxo-dG like Sod2 [40], dOgg1 and Ribosomal protein S3 (RpS3) [41], or "DNA-(apurinic or apyrimidinic site) lyase activity", we hypothesize that the observed increase in C>A transversion may derive from tumour-type specific differences in metabolic activity and the consequent changes in oxidative stress levels.

The SNPs found in our cohort are scattered over the chromosomes and, unlike CNVs, they are not more frequent in the X chromosome than in the autosomes (Fig. 4A, C). The lower rate of mean SNPs/Mb in all chromosomes in female samples may simply reflect the fact that the bratL1 and bratL2 lines, which present the lowest incidence of SNPs, are female and account for most (5/7) of the female samples of the cohort. By analysing groups of SNPs separated by at most 50Kb we identified 96 regions where SNPs appear to be significantly (p≤0.001) clustered in each tumour line (Table S4). However, none of our tumour samples showed any evidence of a “mutator phenotype” following [42]. The longest consecutive series of such clusters (about 400 Kb) maps to a chromosomal region that presents overall enrichment of SNPs, and that spans 3Mb in 3R. 17% (23/134) and 15% (22/151) of the SNPs found in the mbtL2 T10A and T10B lines, respectively, fall within this region, a highly significantly (p≤1x10^−12^) increase compared to the 2% expected if SNPs were randomly distributed along the third chromosome.

### SNPs rates in Drosophila brain tumours are within the range reported for human tumours

To compare the rate of SNPs in our tumour cohort to those reported for human tumours [43] we determined the frequency of the various types of SNPs classified by their localisation in the corresponding gene and deduced the rate of SNPs per Mb in the fraction of the exome that is sufficiently covered for significant SNP calling, considering only those SNPs with a minimum alternative allele frequency of 0.1 (Table S6). For tumour lines with more than 100 SNPs, the fraction of SNPs falling in the exome ranges between 15 and 34% of which more than 60% affect protein sequence. The corresponding percentages are not significant in the lines that present fewer than 100 SNPs. Mean SNPs rate in *brat* tumours (0.16 SNP/Mb of exome) is close to that of the human tumours with the lowest rate of SNPs, like rhabdoid tumour (Fig. 5). Mean SNPs rate in *l(3)mbt* tumours (0.48 SNP/Mb of exome) is within the range of pediatric medulloblastoma and neuroblastoma. Finally, the rates of SNPs in the exome in *aurA* and *lgl* (3.6 SNP/Mb of exome) fall among those of human tumours with a medium-high rate of SNPs, like glioblastoma multiforme, and colorectal cancers (Fig. 5).

**Fig. 5.**
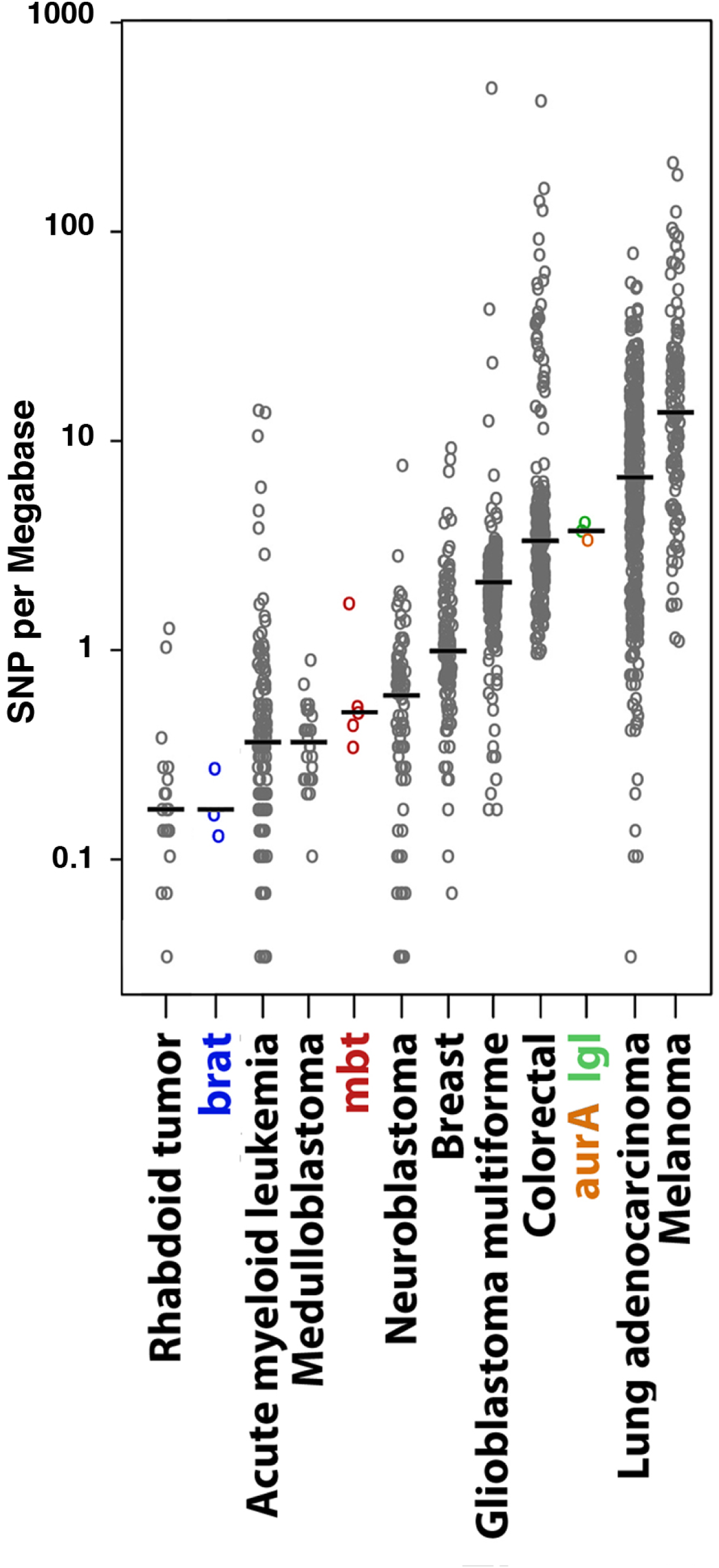
SNP rates of Drosophila larval brain tumours compared to the SNP rate spectrum of a selection of human cancers. Scattered plot of the rates of SNPs/Mb of exome found in late stages (T5 and T10) of the Drosophila cohort (coloured) together with those from a selection of human cancer samples (grey circles; modified after [43].

Malignancy traits are known to worsen over time in the tumours of our cohort: the later the round of implantation the higher the percentage of allografts that develop as tumours, and the shorter the life expectancy of implanted hosts [29, 30]. This observation strongly suggest the acquisition of driver mutations as tumours age. Such is the case in many human cancer types [9] [44] as well as in established Drosophila cell lines which acquire pro-proliferation and anti-apoptotic mutations [27]. However, we have found no genes mutated in more than one tumour line, not even among those with the highest rates of SNPs. Moreover, the fraction of SNPs that are passed on to later time points is very small ranging between 9 and 24 from T0 to T5 and between 0 and 8 from T5 to T10 (Table S6). Thus, for instance, only 3% of the 476 SNPs found in lgl T5 were passed on to lgl T10. Altogether, these results do not support the presence of driver mutations in the cohort that we have analysed. The point has to be made, however, that for detection of driver genes in human cancer, sample sizes are much larger than ours, in the order of hundreds per tumour type [45]. Therefore, the fact that our data does not reveal driver mutations in our cohort of Drosophila larval brain tumours does not rule out their existence.

In summary, we have found that Drosophila larval brain malignant neoplasms with diverse origin present different SNP burdens that are well within the range of SNPs rates reported for human cancer. The very low percentage of SNPs passed on to later time points and the absence of genes mutated in more than one line strongly argues that, like CNVs, tumour-linked SNPs are passenger mutant. The very predominant transvections are likely to result from enhanced oxidative stress conditions that are linked to tumour growth.

## MATERIALS AND METHODS

### Fly strains

All fly stocks and crosses were maintained in standard food medium at 25°C unless otherwise specified. Flies carrying the following mutants and transgenes were used: *pUbiGFP–tub84B* and *pUbi-His2Av::EYFP* [46] *l(3)mbt*^ts1^ [47], *brat*^k06028^ [48], *aurA*^8839^ [16], *l(2)gl*^4^ [19]. The genotypes of each of the tumour lines are as follows. Lines mbtL1 and mbtL2: *Df(1)y-ac w*^*1118*^*, pUbi-His2Av::EYFP, pUbq-alpha-tub-84::GFP*; *l(3)mbt*^ts^1^.^ Lines bratL1 and bratL2*: P{w*^*+*^*, lacW}brat*^*k06028*^ (on a w^+^ background). Line *lgl*: *l(2)gl*^*4*^ Line *aurA*: *w*^*1118*^*, pUbi-His2Av::EYFP, pUbq-alpha-tub-84::GFP; aurA*^8839^. To generate *l(3)mbt* tumour larvae were raised at 29°C.

### Allografts and DNA isolations

Allografts were performed as previously described [7] with minor modifications. Single optic lobes from 3^rd^ instar larvae were dissected and injected into the abdomen of *w*^*1118*^ adult females. Flies were monitored daily and tumours were dissected out when they filled the abdomen of the host. Dissected tumours were resuspended in 100μl of PBS. An aliquot of 5μl of the tumour cell suspension was re-implanted in a new host and the remaining 95μl were processed for DNA isolation by standard lysis-ethanol precipitation, RNAse treatment, and beads-purification (Agencourt AMPure XP, Beckman Coulter). DNA from non-tumour larval tissues was isolated following the same protocol.

### Pair-end DNA sequencing

DNA samples from tumours and their relative controls were processed in parallel. Genomic DNA of each sample was extracted and then fragmented randomly by sonication. After electrophoresis, DNA fragments of about 150-300 bp were purified. Adapter ligation and DNA cluster preparation are performed by Illumina Nextera DNA Sample Preparation KIT (Illumina), and tumour and non-tumoural controls were sequenced in parallel by Illumina Hiseq2000. We performed read quality control using the FastQC software (http://www.bioinformatics.babraham.ac.uk/projects/fastqc). All samples passed minimum quality requirements.

### Alignment and coverage computation and correction (for CNV analyses)

100 bp paired end reads were aligned to the dm6 Drosophila genome version using the STAR aligner [49] with default parameters. Each chromosome was binned into 1000bp segments for which mean coverage was computed using the IGVtools software [50]. We detected uneven coverage for regions with different GC content levels. In order to correct for this bias we fitted a generalized linear model using the Tweedie family with parameter 1.5 and log link function as implemented in the “gam” function from the R statistical language package “mgcv”. Residuals were used for all subsequent calculations.

### Filtering, normalization, segmentation and CNV calling

We downloaded mappability information for the dm3 genome version from and converted coordinates to the dm6 version using the liftOver tool in [51]. Mean GC content was computed for each 1kb bin from the dm6 genome version.

Bins with mappability values of 0 and GC content below the lower .08 quantile were removed from the analysis. Corrected and filtered coverage was quantile normalized for all samples using the function “normalize.quantiles” from the “preprocessCore” R package. Genome segmentation was performed according to [52] using the “segment” function as implemented in the “DNAcopy” R package “CGHcall”. Segmentation and all subsequent steps were performed for each tumour type independently. For each comparison of interest, the ratio was computed between the sample and its corresponding control. Ratios were further normalized using the “normalize” function from the same package. p-value cutoff was set to 0.01 and a minimum of 3 standard deviations between segments. Segment means were normalized using the “postsegnormalize” function. We used the “CGHcall” function from the “CGHcall” package to classify segments into double deletion, single deletion, diploid, single amplification and high amplification. Default parameters were used throughout the analysis.

Gene annotation was performed using the “biomaRt” R package [53] version from May 2015.

In order to obtain CNVs outside of URs we removed bins overlapping with these regions and repeated the segmentation step and CNV calling. Repeat masker regions were downloaded from UCSC for Drosophila melanogaster dm6 version. Under-replicated regions were obtained from [23].

### Alignment and read processing for SNP calling

Reads were aligned to the dm6 version of the Drosophila genome using the BWA software version 0.7.6A [54] with default parameters. The resulting output was converted to the bam format and sorted using samtools version 0.1.19 [55]. We then proceeded to process the data with the software package GATK version 2.5-2 [56] according to their recommended best practices and with default parameters. We used a database of known SNPs downloaded from http://e68.ensembl.org/Drosophila_melanogaster corresponding to the dm3 genome version and converted to the dm6 version using the liftOver tool. Each sample was pre-processed according to the following steps: removal of duplicates using picard version 1.92; realignment of reads around indels using the GATK package with functions RealignerTargetCreator and IndelRealigner; base recalibration with the SNP database mentioned above and the function BaseRecalibrator from GATK.

### Somatic mutation calling

Preprocessed files were used as input for the muTect software version 1.1.4 [57] with default parameters. Each sample was paired with its corresponding control. Resulting somatic SNPs were annotated using the software SNPeff version 3.0 [58].

### SNP clustering

We counted the number of SNPs in windows of 50KB around each SNP detected by our method. We then performed a binomial test assuming a constant probability of finding a SNP in every position of the genome. The total effective size was computed as the number of positions with sufficient information in order to call a SNP. The Mutect algorithm internally defines these positions.

### Gene Ontology Enrichment

Gene ontology enrichment analysis was performed at the Gene Onto Consortium Website (http://geneontology.org) querying the set of 1791 genes that are amplified in at least 1 sample and never deleted and the set of 1101 genes that are deleted in at least 1 sample and never amplified in our cohort. We compared our gene set to the GO cellular component and biological function complete Data Sets and using the Bonferroni correction for multiple testing.

## Statistical Analyses

Unless otherwise stated all statistical test in this study were calculated by the Mann-Whitney test.

## ACKNOWLEDGEMENTS

We thank the Bloomington Drosophila Stock Centre for providing fly stocks and A. Duran for technical assistance. Research in our laboratory is supported by ERC AdG 2011 294603 Advanced Grant from the European Research Council; BFU2015-66304-P and Redes de Excelencia BFU2014-52125-REDT-CellSYS from the Spanish MINECO, Spain; and SGR Agaur 2014 100 from Generalitat de Catalunya, Spain.

## SUPPORTING INFORMATION LEGENDS

**Fig. S1.**
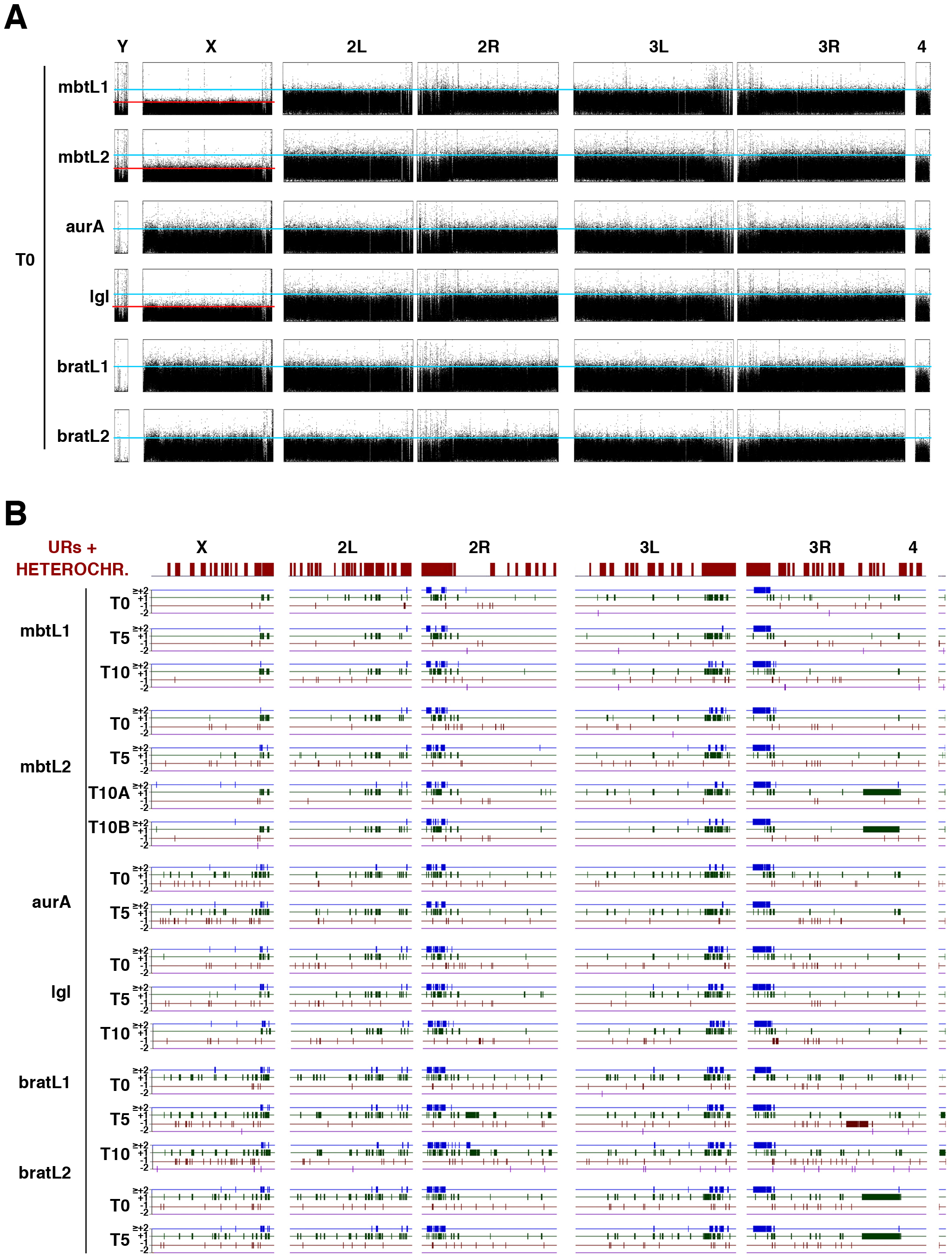
Sequence coverage and first draft map of CNVs. A) Overview of sequence coverage over the genome at the first round of allograft (T0). The halved coverage of the X chromosome compared to that of the autosome arms and the significant coverage of Y chromosome specific sequences in mtbtL1, mbtL2, and lgl indicates that these lines originated in male larvae. B) Map of CNVs identified in different lines at different time points. Copy number gains (≥+2, blue; +1, green) and losses (-1, red; and ≤-2, purple) are mapped along chromosome arms X, 2L, 2R, 3L, 3R, and 4th. The heterochromatic Y chromosome is omitted. A very significant fraction of copy number gains map on under-replicated regions (URs and heterochromatin; shown in brown at the top of the map).

**Table S1.**

Catalogue of CNVs found in the cohort.

**Table S2.**

GO analyses of genes affected by CNVs.

**Table S3.**

Catalogue of SNPs found in the cohort.

**Table S4.**

SNP cluster analyses.

**Table S5.**

SNPs types found in the cohort.

**Table S6.**

Percentage SNPs passed on to later time points.

